# Successfully treating biofilms of extensively drug resistant *Klebsiella pneumoniae* isolates from hospital patients with N-Acetyl Cysteine

**DOI:** 10.1101/2022.09.07.506922

**Authors:** Ankurita Bhowmik, Sambuddha Chakraborty, Anusha Rohit, Ashwini Chauhan

## Abstract

*Klebsiella pneumoniae* is one of the leading causes of community and nosocomial infections. Reduced treatment options against extensively drug resistant (XDR) - *K. pneumoniae,* is a serious concern in hospital settings, and hence, WHO has categorized it as a “critical priority pathogen”. Biofilm forming ability is a common virulence mechanism amongst *K. pneumoniae* that is associated with antibiotic tolerance up to 1000X MIC and hence, are difficult to treat. N-acetyl cysteine (NAC) is an FDA approved mucolytic drug used to treat acetaminophen-associated toxicity and obstructive pulmonary diseases. In this study, we assessed NAC’s antibacterial and antibiofilm activity against clinical isolates of XDR *K. pneumoniae,* obtained from Madras Medical Mission Hospital, India. To assess the biofilm eradication ability of NAC, we grew biofilms in 96 well plates and treated the mature biofilms with different concentrations of NAC. We observed that the biofilms of only 3 isolates of XDR *K. pneumoniae* could be eradicated at a concentration as low as 20mg/ml. Although increasing the concentration of NAC to 80mg/mL could significantly reduce the biofilms of all the isolates up to 4-5 Log, NAC at a concentration of 100 mg/mL successfully eradicated the mature biofilms of all the isolates of XDR *K. pneumoniae*. This *in vitro* study demonstrates the potential of NAC as an efficient agent against the biofilms of clinical isolates of XDR-*K. pneumoniae* and thus, provides a promising alternative to antibiotics.

## 1. Introduction

*Klebsiella pneumoniae* is a ubiquitous, Gram negative, non-motile, and encapsulated bacillus that colonizes skin, mouth, respiratory tract, and digestive tract in human. It is a major cause of community and hospital acquired pneumonia, urinary tract infection, sepsis, and meningitis in immunocompetent and immunocompromised patients (Joseph et al., 2021; Liu et al., 2021). Recent studies have demonstrated the association of *K. pneumoniae* with severe infections including pyogenic liver abscesses; endophthalmitis etc. (Russo & Marr, 2019). Diverse virulence factors such as siderophores, capsular polysaccharides, lipopolysaccharides, and fimbrial adhesins contribute to the pathogenicity of *K. pneumoniae* (Remya et al., 2019; Gomes et al., 2021). Moreover, the propensity of *K. pneumoniae* to acquire multidrug resistance has made them more notorious to deal with in the hospital environment. In 2020, the Indian Council of Medical Research (ICMR) categorized *K. pneumoniae* as a Group-I pathogen due to the emergence of resistance toward the last line of antibiotics resulting in increased morbidity and mortality rate. The global priority pathogen (GPP) list published by WHO in 2017 has also enlisted ESBL producing carbapenem-resistant *K.pneumoniae* among the “critical priority pathogens” (WHO 2017). Moreover, *K. pneumoniae* was recently reported to be prevalent in up to 18% of clinical specimens analyzed in India. Furthermore, ICMR has also disclosed an extended resistance in *K. pneumoniae clinical* isolates towards cephalosporins, aminoglycosides, carbapenem, fluoroquinolones, and polymyxin *K. pneumoniae* (Indian Council of Medical Research, 2020; Ashwath et al., 2022). *K. pneumoniae* possesses a natural resistance to the penicillin class of antibiotics due to the production of sulfhydryl variable (SHV-1) penicillinases (Kakoullis et al., 2021). In addition, the presence of plasmid encoded extended spectrum beta-lactamases (ESBL) including cefotaximase-M (CTM-M) in *K. pneumoniae* confers resistance towards broad spectrum aminoglycosides and cephalosporins (Piperaki et al., 2017; Karaiskos & Giamarellou, 2020). Moreover, different carbapenemases such as metallo-beta-lactamases, *Klebsiella pneumoniae* carbapenemases, and oxacillinases confers carbapenem resistance in *K. pneumoniae* (Lopes et al., 2020).

Besides different resistance mechanisms, biofilm forming ability of *K. pneumoniae* confers additional protection from antibiotics and the host immune system (Guilhen et al., 2019; Singh et al., 2021). The biofilm matrix enables the development of heterogenous bacterial subpopulations including metabolic inactive persister cells due to gradients of oxygen, nutrients, metabolic products, and accumulated toxins. The persister cells can survive lethal dosage of antibiotics and hence, confer increased antibiotic tolerance (Chauhan et al., 2014; Lebeaux et al., 2014). Biofilm development on indwelling devices and epithelial tissues is clinically significant and is widely reported (Chauhan, et al., 2012; Lebeaux et al., 2013).

Globally, the rise and spread of XDR-*K. pneumoniae* infections have renewed the research interest in studying the correlation between antibiotic resistance and biofilm encoding genes in *K. pneumoniae* (Effah et al., 2020; Ballén et al., 2021; Shadkam et al., 2021). Currently, different combinations of antibiotics such as meropenem with fosfomycin, and/or gentamicin, ceftazidime/avibactam with fosfomycin, or gentamicin or tigecycline, etc. are used to treat XDR-*K. pneumonia* infections (Daikos et al 2014; Jacobs et al 2017; Karaiskos et al 2019; Petrosillo et al 2019; Yu et al 2019; Bassetti et al 2021). Besides, alternatives to antibiotics have also been explored such as bacteriophage therapy in conjunction with non-active antibiotics such as sulfamethoxazole-trimethoprim was effective against XDR-*K. pneumoniae* isolate (Bao et al., 2020). Recent reports demonstrated the potential of antimicrobial peptides such as AA139, SET-M33, and ΔM2 against colistin resistant *K. pneumoniae* (van der Weide et al., 2019; Rivera-Sánchez et al., 2020). EDTA in combination with colistin is recently reported to manage colistin resistant *K. pneumoniae* (Shein et al., 2021). However, to manage the drug resistance challenge among *K. pneumoniae,* constant exploration of novel non-antibiotic therapeutic interventions is urgently required.

N-acetyl cysteine (NAC) is a synthetic derivative of endogenous amino acid, L-cysteine, and a precursor of glutathione. It is an FDA approved mucolytic drug used for decades to treat acetaminophen overdose and obstructive pulmonary diseases (Tenório et al., 2021). Due to its mucolytic activity, NAC has attracted microbiologists to explore its potential as an antibacterial and antibiofilm agent. A number of in vitro studies assessed the activity of NAC against different Gram positive and Gram-negative bacteria including *Escherichia coli, Acinetobacter baumannii, Pseudomonas aeruginosa, Klebsiella pneumoniae, Staphylococcus aureus, and Staphylococcus epidermidis* amongst others (Quah et al.,2012; Choi et al.,2018; Feng et al.,2018; Jun et al.,2019; Ciacci et al., 2019; Li et al.,2020; Guerini et al.,2020; Kundukad et al., 2020; Manoharan et al., 2021). Given the broad-spectrum activity of NAC against bacteria, this study aims at repurposing N-acetyl cysteine (NAC) to treat biofilms of XDR *K. pneumoniae,* isolated from patients in hospitals.

In our study, a total of 1402 Gram-negative clinical isolates were collected during the period from 2018 to 2020 at Madras Medical Mission Hospital, Chennai India. Out of this, 374 (26.6%) were identified as *Klebsiella spp*. and intriguingly, 99% of isolates amongst the *Klebsiella spp*. were *K. pneumoniae*. Based on the prevalence and resistance profile, we selected 12 XDR *K. pneumoniae* isolates and assessed the *in vitro* antibacterial as well as antibiofilm activity of NAC against these isolates. We observed that NAC was bactericidal against all the XDR *K. pneumoniae* isolates at a concentration of 10mg/mL. Furthermore, NAC could successfully eradicate *in vitro* biofilms of all the XDR *K. pneumonia* at an acceptable concentration of 100mg/mL (Olsson et al., 1988; Hughes et al., 2018; NIH U.S. National Library of Medicine 2018, Guerini et al., 2022). Safe alternatives to antibiotics are a compelling need of the hour to treat *K. pneumoniae* infections to improve patient outcomes. N-acetyl cysteine may be a safe option for treating XDR *K. pneumoniae* leading to reduced socio-medico-economic burden.

## 2. Materials and Methods

### 2.1. Bacterial strains and Growth media

All the *XDR-Klebsiella pneumoniae* isolates were grown at 37°C in Luria-Bertani Broth (LB) purchased from Himedia, India (M1245). For viable cell estimation the serially diluted culture was plated on sterile LB agar plates and incubated overnight at 37°C.

### 2.2. Antimicrobial agents

N-acetyl cysteine (NAC) was purchased from Sigma Aldrich (A7250). Isotonic NAC solution was prepared by dissolving NAC in sterile 0.9% saline.

### 2.3. Estimation of antibiotic susceptibility

VITEK 2 compact (bioMerieux, France) was used for biochemical characterization and determining the antibiogram of all the XDR-*K. pneumoniae* isolates using disposable antibiotic susceptibility Testing (AST) cards. VITEK 2 instrument continuously monitors the bacterial growth and records the fluorescence, turbidity, and photometric signals in presence of an increasing concentration of various antibiotics. Wells with no antibiotics served as the positive control (Ligozzi et al 2002).

### 2.4. Assessing the Minimum inhibitory Concentration (MIC)

The MIC of NAC against the clinical isolates of XDR-*K. pneumoniae* was determined by broth microdilution assay (Chauhan et al., 2012; Lebeaux et al., 2014; Lewis et al.,2022). Briefly, NAC was serially diluted in LB in a microtiter plate to give a 2-fold serial dilution. An equal volume of exponentially growing *K. pneumoniae isolate* was added to give a final inoculum of 5 × 10^5^ CFU/mL in all the wells except the negative control. Well with bacteria but no NAC, served as an untreated control. The plate was incubated for 18h at 37°C under static conditions and the first well without visible growth was noted as the MIC concentration. All the experiments were done at least in triplicate.

### 2.5. Effect of NAC on Exponential phase planktonic XDR-*K. pneumoniae*

Growth inhibition efficacy of NAC against clinical isolates of XDR-*K. pneumoniae* was done in 96 well microtitre plates using a BioteK Epoch-2 microtiter plate reader (Agilent Technologies). Each treatment was replicated in at least 3 wells. Log phase culture (OD600 ≈ 0.3-0.5) of *K. pneumoniae* isolates were treated with sub-MIC (2.5 or 5mg/mL) concentration to 80mg/mL of NAC for 24h at 37°C with continuous shaking. Untreated wells served as control. The OD600nm were measured by the Biotek Epoch-2 plate reader for 24 hrs. After 24h, the viable cell count was estimated by plating the serial dilutions on LB agar media, and plates were incubated at 37°C overnight (Lebeaux et al., 2014).

### 2.6. *In vitro* biofilm forming ability of Clinical isolates of XDR *K. pneumoniae*

The biofilm forming ability of the clinical isolates of XDR *K. pneumoniae* was determined by assessing the ability of the cells to adhere to the wells of 96-well polystyrene microtiter plates by previously described crystal violet (CV) staining method (O’Toole et al.,1998; Chauhan et al 2013). An overnight culture of *K. pneumoniae* grown in LB at 37°C under shaking was diluted 1/100 in fresh LB (1uL inoculum in 100uL LB media per well) and incubated at 37°C under static conditions for 72h. After incubation, the non-adherent cells were gently removed. The wells were gently washed thrice with 0.9% saline and 125uL 0.1% CV stain was added to the wells. The plates were incubated at room temperature for 15 mins and the wells were rinsed with 0.9% saline. The plates were then air dried for 15mins before solubilizing the CV with acetone-ethanol (1:4), and incubated at room temperature for 15 minutes. Biofilm biomass was estimated as a function of CV stain by measuring the absorbance at 595nm. The results are an average of five technical replicate wells from three independent experiments. The wells containing media without bacteria were used as the negative control, and strain *K. pneumoniae* KPP1 and KPP2 were used as positive controls as they form strong biofilms under invitro laboratory conditions (unpublished data). The cut-off OD (ODc) is defined as three standard deviations (SD) above the mean OD of the negative control, i.e ODc= average OD of negative control+ (3xSD of negative control). XDR-*K. pneumoniae* isolates were categorized as negative, weak biofilm formers, moderate or strong biofilm formers as follows: negative biofilm formers: OD ≤ ODc; weak biofilm producer: (OD_c_ < OD ≤2× OD_c_), moderate biofilm producer (2× OD_c_ < OD ≤4× OD_c_), strong biofilm producer (OD >4× OD_c_).

### 2.7. *In vitro* assessment of the inhibitory potential of NAC against the biofilms formed by Clinical isolates of XDR *K. pneumoniae*

The biofilm inhibition assays were done as described by Lebeaux et al 2014 with slight modifications (Lebeaux et al., 2014). The overnight cultures of *K. pneumoniae* isolates were inoculated in fresh media at 1:100 dilution in a 96-well microtiter plate and incubated for 96h to form mature biofilm at 37°C without shaking. The planktonic bacteria were removed by gentle pipetting and the wells were rinsed thrice with 0.9% saline. The 96h old mature biofilms growing in the micro-titre plates were treated with different concentrations of isotonic solution of NAC (5mg/mL, 10mg/mL, 20mg/mL, 40mg/mL, 80mg/mL and 100mg/mL) by directly adding 125uL NAC solution into the wells. The treatment was allowed to dwell in the wells for 24h at 37°C under static conditions. 0.9% saline treated wells served as untreated controls. After 24h, the NAC solution was removed from the wells and rinsed thrice with 0.9% saline. Wells were scraped vigorously to remove any residual biofilm and homogenized with 0.9% saline. The viable cell count was estimated by plating the serial dilutions on LB agar medium and the plates were incubated at 37°C. Each treatment was given in replicate of four and each experiment was repeated at least thrice.

### 2.8. Cytotoxicity assay

Cytotoxicity of N-acetylcysteine (NAC) was assessed on L929 (mouse fibroblasts), and J774A.1 (mouse macrophages) cell lines (procured from National Centre for Cell Sciences, Pune, India). Two concentrations of NAC; 80 mg/mL & 40 mg/mL in 0.9% isotonic saline solution were subjected to the test. The assay was performed using CyQUANT™ MTT Cell proliferation assay kit, Invitrogen. Briefly, approximately 4000 cells of both cell types were seeded per well of 96-well microtiter plates and kept for incubation at 37 °C with 5% CO2. After attainment of 70-80% confluency, spent media were discarded from all the wells including the control wells. Two test concentrations of NAC were inoculated in the designated wells. Control wells were left with the media. The plates were incubated at 37 °C with 5% CO2 for 30 minutes. Post incubation, NAC was aspirated out from every test well and replaced by fresh media. MTT [3-(4,5-dimethylthiazol-2-yl)-2,5-diphenyl tetrazolium bromide] solution was added to all the treated and control wells, and the plate was incubated at 37 °C for four hrs. in 5% CO2. Post MTT incubation, all the contents from each well were removed, followed by the addition of DMSO. After adding DMSO, the plates were incubated for 10 minutes, and the absorbance was measured using EPOC 2, 96 well plate reader (BioTek) at 570 nm.

### 2.9. Statistical analysis

All the results are replicates of at least three independent experiments. Statistical differences were evaluated using one-way analysis of variance (ANOVA) using Graphpad Prism version 8.0. *P* values lower than 0.05, were considered statistically significant.

## 3. Results

### 3.1. *K. pneumoniae* is most prevalent amongst the Gram negative bacteria isolated from clinical settings

A total of 1402 Gram-negative bacterial samples were isolated from patients admitted to Madras Medical Mission Hospital during 2018-2020. Among these, *Klebsiella spp*., was found to be the second predominant bacterial species among all the Gram-negative bacteria. *Klebsiella spp*., comprised 26.6% of the isolated Gram-negative bacteria. Intriguingly, 99% of isolates amongst the *Klebsiella spp*. were *K. pneumoniae* and only 1% was *K. oxytoca* (Fig.1).

**Fig 1:**
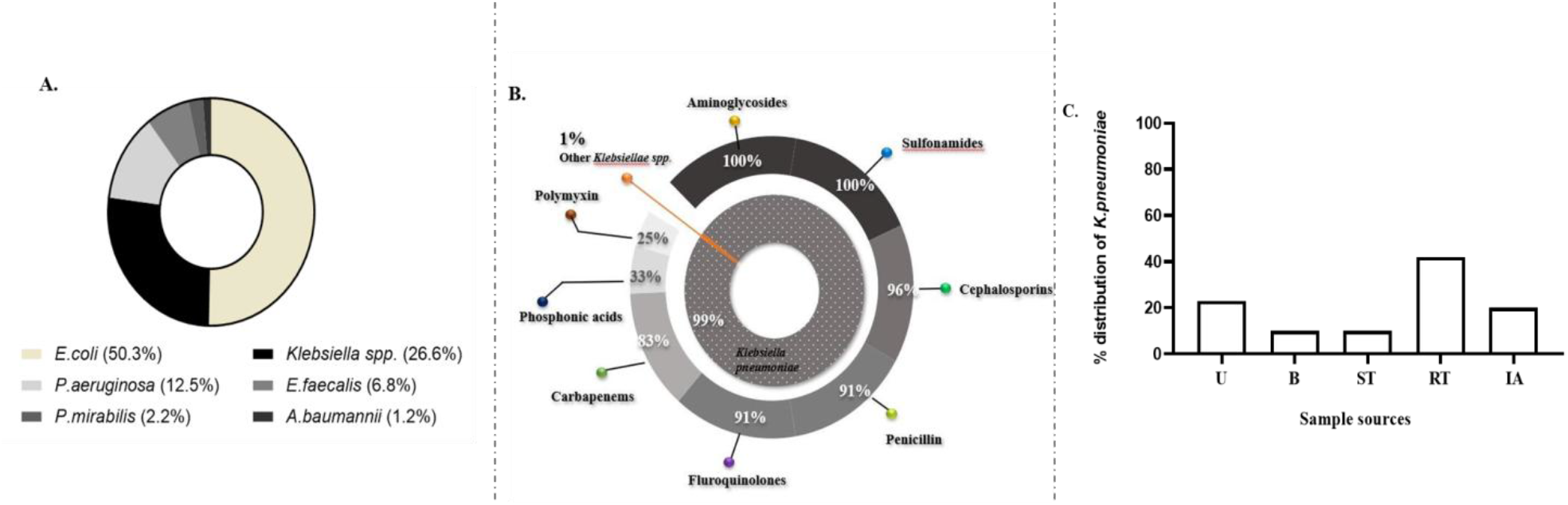
Prevalence of *Klebsiella spp.* and its drug resistance profile in clinical settings of Madras Medical Mission Hospital, India. A total of 1402 Gram negative bacteria were isolated from hospital settings and characterized for identification. **A.** *Klebsiella spp*. comprised 26.6%, amongst the isolated Gram negative bacteria. **B.** Of all the *Klebsiella spp*., 99% were *K. pneumoniae* that showed resistance to multiple classes of antibiotics including carbapenems and polymyxin. **C.** *K. pneumoniae* was more prevalent among samples isolated from patients with respiratory tract infections (42%). U-Urine, B-Blood, ST-Soft tissues, RT-Respiratory tract, IA-Intraabdominal.

### 3.2. *K. pneumoniae* isolates showed resistance to the different classes of antibiotics

The characterization and antibiotic susceptibility profiling of the *K. pneumoniae* isolates was done by an automated Vitek 2 compact system (bioMerieux, France). All the *K. pneumoniae* isolates showed resistance to multiple classes of antibiotics (Fig 1B). We selected 12 XDR-*K. pneumoniae* isolates in our study as they showed resistance to almost all the tested antibiotics (Fig 2). It is worrisome to note that all the isolates showed 100% resistance towards amikacin, amoxicillin-clavulanate, cefazolin, gentamicin, ceftriaxone, cefuroxime, and co-trimoxazole. Furthermore, only less than 20% of isolates were susceptible to imipenem, meropenem, ertapenem, pip-tazobactam, and cefoperazone sulbactam. Most of the isolates (90%) showed resistance towards tigecycline, netilmicin, ciprofloxacin, ofloxacin, and cefepime. Colistin is currently used as a final resort for treating multidrug resistant *K. pneumoniae*. However, genetic mutations in the *phoPQ* and *pmrAB* system result in colistin resistance in *K. pneumoniae* (Nirwan et al., 2021). Recent reports also suggest *mcr* plasmid-mediated colistin resistance in *K. pneumoniae* (Karki et al., 2021; Salloum et al., 2020). In this study, we also observed that 25%*K. pneumoniae* isolates were colistin resistant, which is concerning, and indicative of the probable spread of colistin resistance genes. A recent report in 2021 showed the tendency of carbapenem resistant *K. pneumoniae* to acquire colistin resistance under tigecycline or colistin pressure (Andrew Evans et al., 2021). Similarly, in our study, we observed that all the colistin resistant *K. pneumoniae* isolates were resistant to carbapenems.

**Fig 2:**
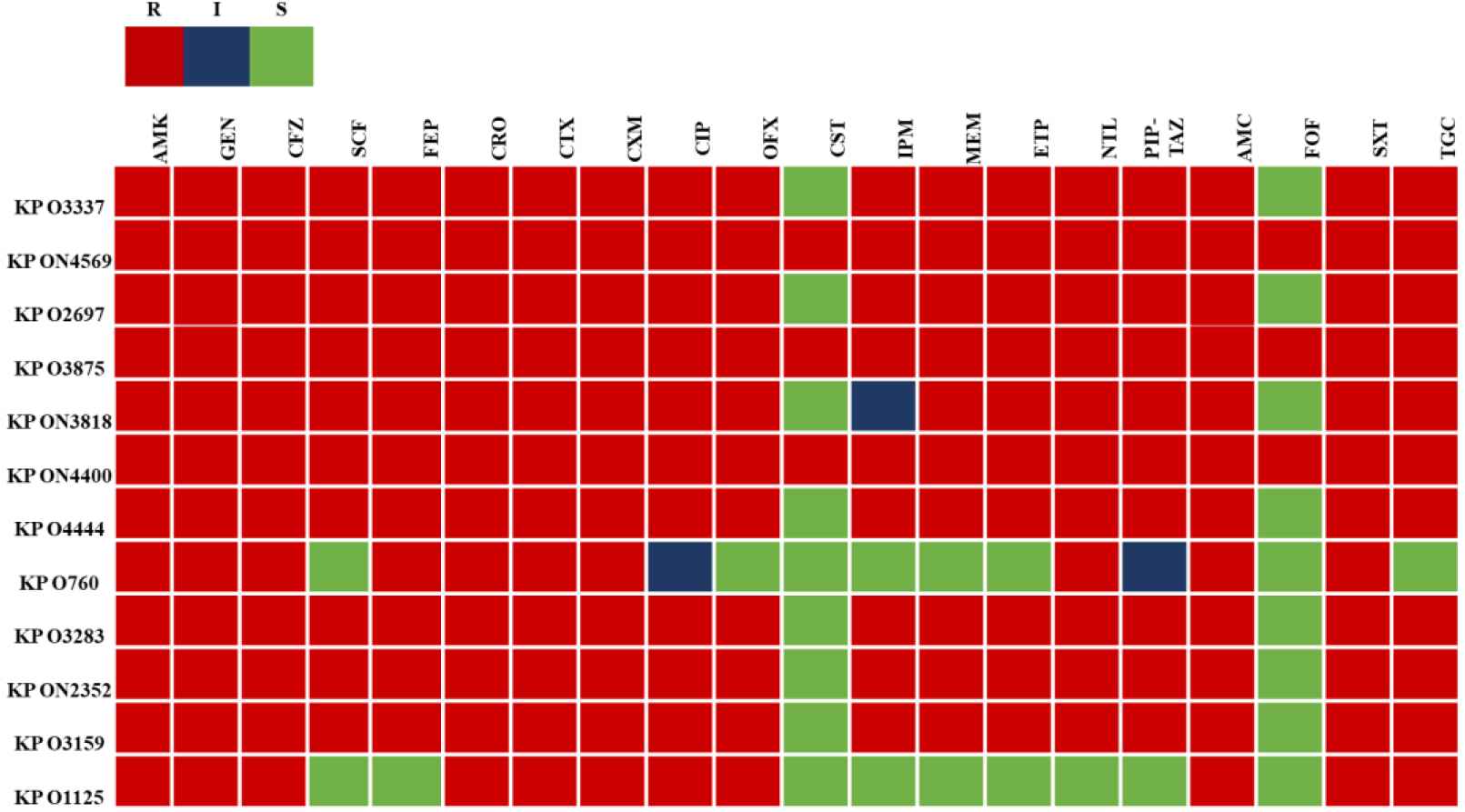
Drug resistance profile of 12 XDR-*K. pneumoniae* isolates used in this study. The heatmap represents the antibiogram and extensive antibiotic resistance among the *K. pneumoniae* isolates. Rows represent *K. pneumoniae* isolates and columns represent different antibiotics. The red blocks signify resistance (R), the blue blocks signify intermediate (I), and the green blocks signify susceptibility (S) to antibiotics. AMK-Amikacin, GEN-Gentamicin, CFZ-Cefazolin, SCF-Cefoperazone sulbactam, FEP-Cefepime, CRO-Ceftriaxone, CTX-Cefotaxime, CXM-Cefuroxime, CIP-Ciprofloxacin, OFX-Ofloxacin, CST-Colistin, IPM-Imipenem, MEM-Meropenem, ETP-Ertapenem, NTL-Netilmycin, PIP-TAZ-Pip tazobactam, AMC-Amoxicillin clavulanate, FOF-Fosfomycin, SXT-Co-trimoxazole, TGC-Tigecycline

### 3.3. NAC effectively kills clinical isolates of XDR-*K. pneumoniae*

To assess the anti-bacterial efficacy of NAC, a weak protic acid, we determined its minimum inhibitory concentration (MIC) against the clinical isolates of XDR-*K. pneumoniae* as per CLSI guidelines. Intriguingly, we observed ~ 83% (10 out of 12) of the XDR *K. pneumoniae* isolates that we chose for the study were susceptible to 5mg/mL of NAC. The remaining 2 XDR *K. pneumoniae* isolates were susceptible to a two-fold more NAC concentration i.e., 10mg/mL (Table 1). Furthermore, we checked the effect of NAC on the cell viability of XDR-*K. pneumoniae* isolates by treating exponentially grown cells with NAC from sub-MIC concentration (2.5 or 5mg/mL) to 80mg/mL for 24h and checked the changes in growth kinetics and colony forming units (CFU) (Table 2) (Fig S.1). We observed that NAC was bactericidal at MIC concentration against all the clinical isolates except KPO760. At MIC (5mg/mL), NAC reduced the growth of KPO760 up to 4.8logs, which completely disappeared with a two-fold increase in MIC (10mg/mL).

**Table 1:** Minimum Inhibitory Concentration of NAC against clinical isolates of XDR *K. pneumoniae*

| Clinical isolates of XDR *K. pneumoniae* (Kp) | MIC (mg/mL) |
| --- | --- |
| ON2352 | 5 |
| O3875 | 5 |
| ON4400 | 5 |
| O3818 | 5 |
| O3337 | 5 |
| O760 | 5 |
| O4444 | 5 |
| O1125 | 10 |
| O3283 | 5 |
| O2040 | 5 |
| ON2697 | 5 |
| O3159 | 10 |
| **\*MIC results are mean results of at least 3 experiments** |  |

**Table 2:**
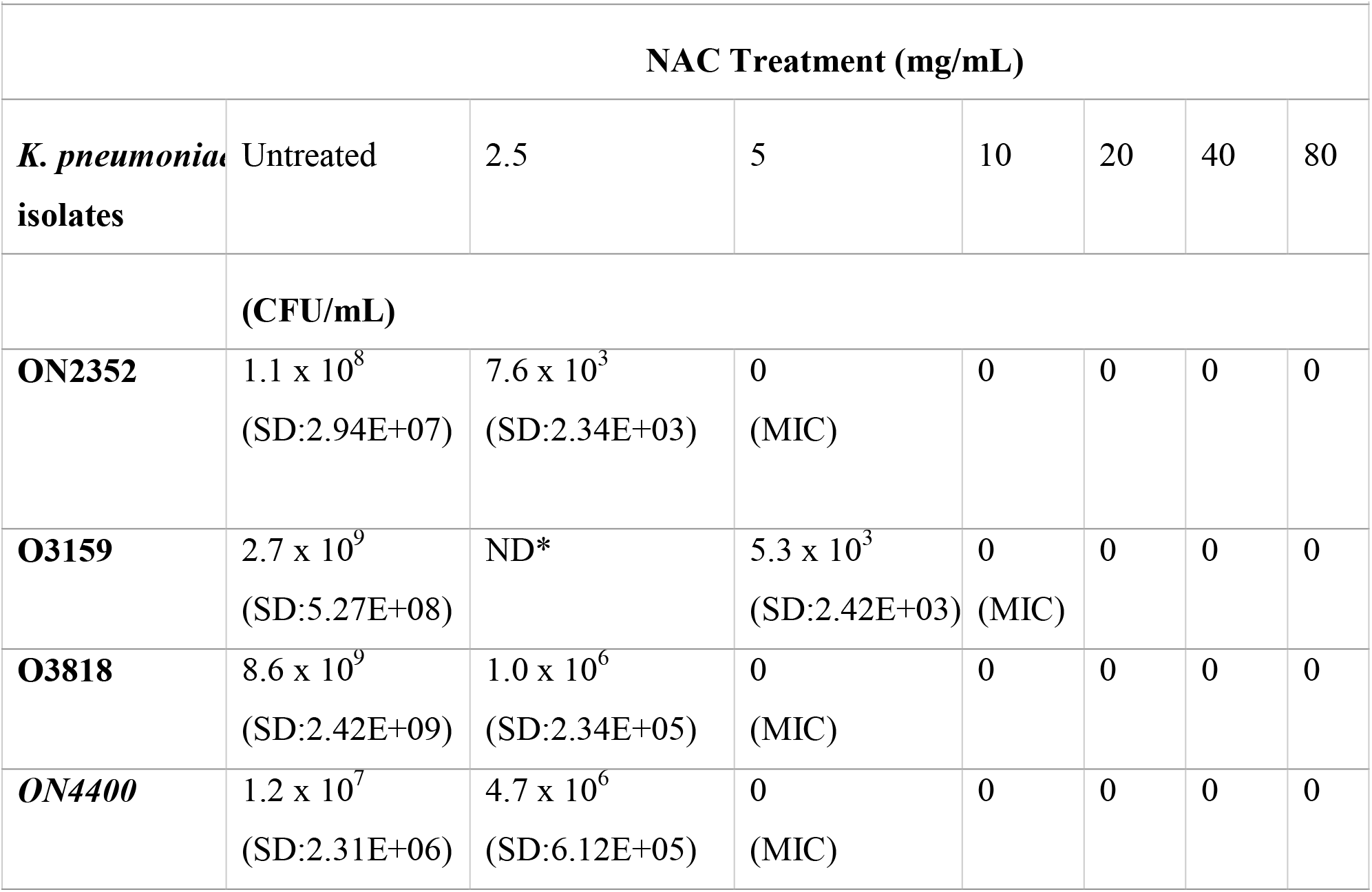
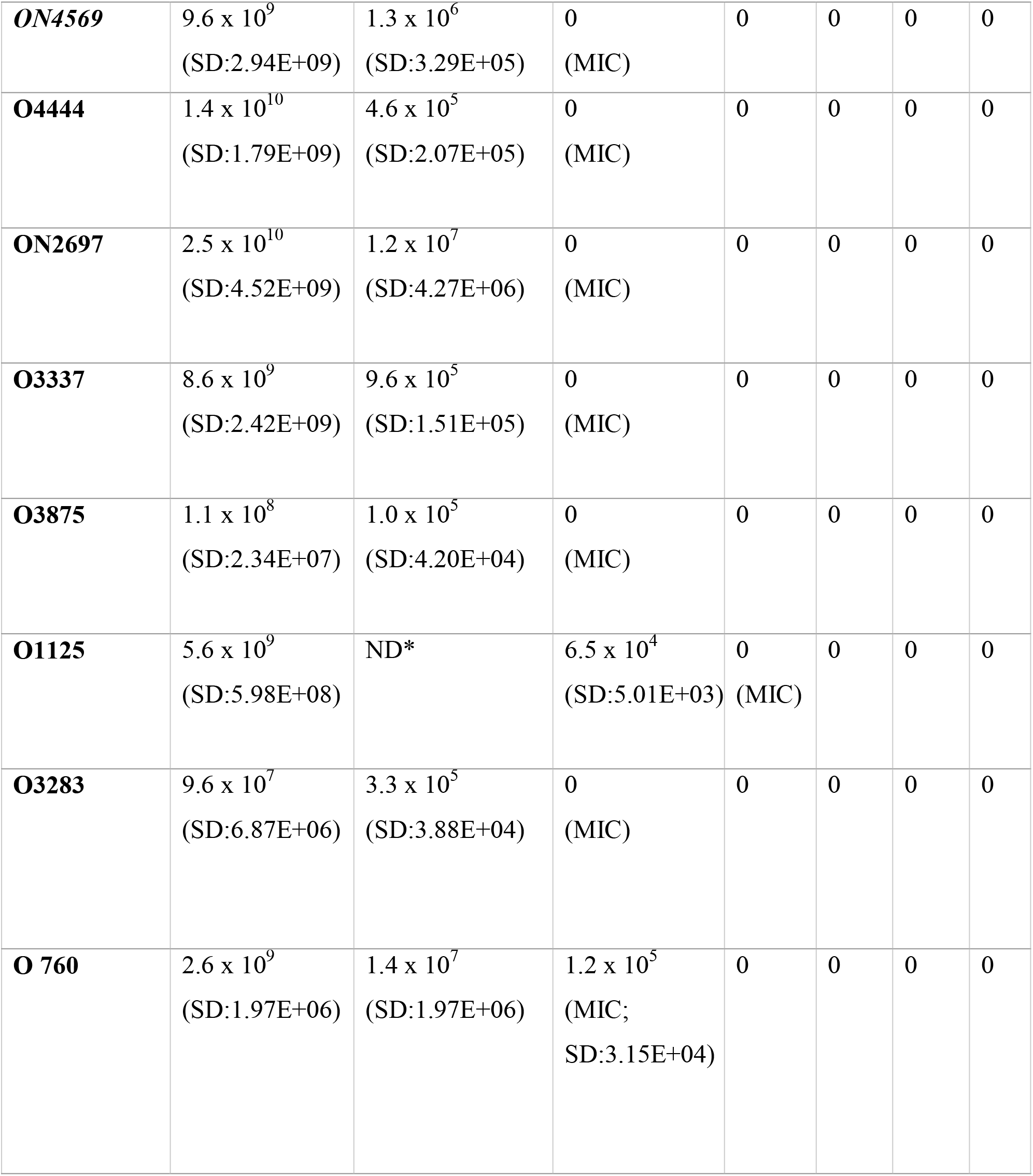
Effect of different concentrations of NAC on exponentially growing clinical isolates of XDR-*K. pneumoniae*. * ND: Not done

### 3.4. Clinical isolates of XDR *K. pneumoniae* are moderate biofilm formers

The biofilm forming ability of 12 XDR-*K. pneumoniae* isolates were assessed by the previously described CV staining method (Stepanović et al., 2007; Leoney et al., 2020). The cut off OD (ODc) obtained was 0.076. We observed that 58% (7/12) of the isolates were moderate biofilm formers, 33% (4/12) were weak biofilm formers and 8% (1/12) were strong biofilm former among the XDR *K. pneumoniae* isolates (Fig.3). Interestingly, three of the *K. pneumoniae* isolates, KPON4569, KPO3875, and KPON4400, that are resistant to all the antibiotics, are weak biofilm formers. All the moderate and strong biofilm formers showed variable antibiotic resistance patterns. It would be interesting to study the effect of multidrug resistance on the fitness of *K. pneumoniae* isolates.

**Fig 3:**
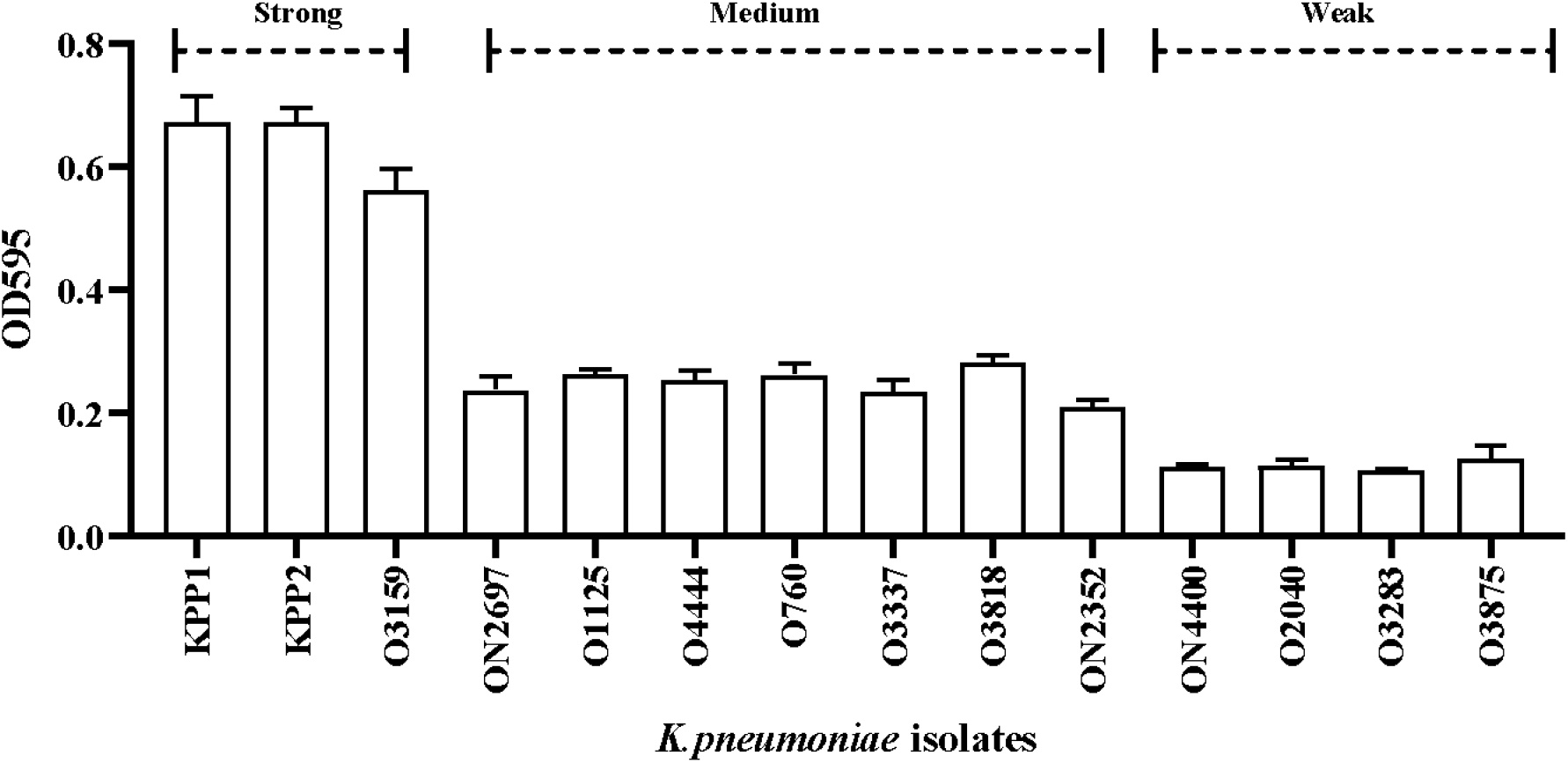
Biofilm producers among XDR-*K. pneumoniae* isolates. The biofilm forming ability of all the isolates was determined by the CV staining method in 96 well plates. The X-axis represents the *K. pneumoniae* isolates and the Y-axis represents the optical density at 595nm. The strains were classified based on their biofilm forming abilities according to previous reports (Stepanović et al., 2007; Leoney et al., 2020).

### 3.5. NAC effectively eliminates mature XDR-*K. pneumoniae* biofilms

Managing biofilm associated infections is challenging especially when the infections are due to antibiotic resistant isolates such as XDR-*K. pneumoniae*. We evaluated the *in vitro* antibiofilm activity of NAC against the 12 XDR-*K. pneumoniae* isolates. The 96h preformed biofilms were treated with different concentrations of NAC prepared in an isotonic solution, ranging from 5mg/mL to 100mg/mL for 24h and the cell viability was estimated by CFU/mL plating. Although we observed a varying level of killing of biofilm-associated XDR-*K. pneumoniae* isolates treated with *K. pneumoniae* NAC upon 24h exposure, NAC could successfully eradicate the biofilms of all the XDR isolates (Table 3).

**Table 3:** Antibiofilm activity NAC against mature biofilm of XDR-*K. pneumoniae* isolates.

|  |  | NAC concentration(mg/mL) |  |  |  |  |  |
| --- | --- | --- | --- | --- | --- | --- | --- |
| **Kp isolates** | Untreated | 5 | 10 | 20 | 40 | 80 | 100 |
| **ON2352** | 2.7 X 10<sup>7</sup><br>(SD:<br>3.27E+06) | 1.9 x 10<sup>5</sup><br>(SD:<br>4.56E+05) | 1.7 x 10<sup>5</sup><br>(SD:<br>8.14E+04) | 4.7 x 10<sup>3</sup><br>(SD:<br>2.42E+04) | 3.3 x 10<sup>3</sup><br>(SD:<br>3.58E+03) | 1.6 x 10<sup>3</sup><br>(SD:<br>8.16E+02) | 0 |
| **O3159** | 1.2 x 10<sup>9</sup><br>(SD:<br>4.27E+08) | 1.1 x 10<sup>7</sup><br>(SD:<br>2.94E+06) | 3.6 x 10<sup>6</sup><br>(SD:<br>2.42E+06) | 1.4 x 10<sup>6</sup><br>(SD:<br>1.41E+06) | 6.3 x 10<sup>4</sup><br>(SD:<br>7.45E+05) | 4.6 x 10<sup>3</sup><br>(SD:<br>3.44E+05) | 0 |
| **O3818** | 1.9 x 10<sup>10</sup><br>(SD:<br>4.34E+08) | 1.3 x 10<sup>9</sup><br>(SD:<br>2.73E+08) | 1.0 x 10<sup>8</sup><br>(SD:<br>1.74E+08) | 9.6 x 10<sup>7</sup><br>(SD:<br>2.10E+07) | 4.3 x 10<sup>6</sup><br>(SD:<br>2.94E+06) | 5.6 x 10<sup>4</sup><br>(SD:<br>2.66E+04) | 0 |
| **ON4400** | 1.0 x 10<sup>10</sup><br>(SD:<br>3.44E+09) | 1.3 x 10<sup>7</sup><br>(SD:<br>3.01E+06) | 1.0 x 10<sup>7</sup><br>(SD:<br>2.53E+06) | 9.0 x 10<sup>6</sup><br>(SD:<br>2.97E+06) | 5.0 x 10<sup>6</sup><br>(SD:<br>2.10E+06) | 4.3 x 10<sup>5</sup><br>(SD:<br>2.34E+05) | 0 |
| **ON4569** | 1.4 x 10<sup>9</sup><br>(SD:<br>4.34E+08) | 7.6 x 10<sup>8</sup><br>(SD:<br>2.42E+08) | 4.6 x 10<sup>8</sup><br>(SD:<br>2.42E+08) | 4.3 x 10<sup>7</sup><br>(SD:<br>2.42E+07) | 4.6 x 10<sup>5</sup><br>(SD:<br>8.91E+05) | 1.1 x 10<sup>4</sup><br>(SD:<br>1.51E+03) | 0 |
| **ON2697** | 9.0 x 10<sup>6</sup> | 7.0 x 10<sup>4</sup> | 0 | 0 | 0 | 0 | 0 |
|  | (SD:<br>2.10E+06) | (SD:<br>3.03E+04) |  |  |  |  |  |
| **O3875** | 6.3 x 10<sup>7</sup><br>(SD:<br>2.8E+07) | 3.3 x 10<sup>5</sup><br>(SD:<br>1.97E+06) | 0 | 0 | 0 | 0 | 0 |
| **O3337** | 1.7 x 10<sup>8</sup><br>(SD:<br>2.42E+08) | 1.0 x10<sup>6</sup><br>(SD:<br>2.07E+06) | 5.6 x10<sup>5</sup><br>(SD:<br>2.97E+04) | 0 | 0 | 0 | 0 |
| **O1125** | 6.3 x 10<sup>8</sup><br>(SD:<br>2.48E+05) | 1.7 x10<sup>7</sup><br>(SD:<br>1.57E+05) | 8.3 x10<sup>6</sup><br>(SD:<br>2.06E+04) | 1.9 x10<sup>5</sup><br>(SD:<br>2.94E+06) | 8.6 x10<sup>4</sup><br>(SD:<br>3.22E+05) | 7.3 x10<sup>3</sup><br>(SD:<br>2.12E+04) | 0 |
| **O760** | 8.0 x 10<sup>9</sup><br>(SD:<br>2.57E+05) | 3.5 x 10<sup>8</sup><br>(SD:<br>3.05E+05) | 1.9 x10<sup>6</sup><br>(SD:<br>1.48E+07) | 7.1 x10<sup>5</sup><br>(SD:<br>4.52E+0) | 1.5 x10<sup>4</sup><br>(SD:<br>2.5E+04) | 4.0 x10<sup>3</sup><br>(SD:<br>1.07E+05) | 0 |
| **O3283** | 9.6 x 10<sup>8</sup><br>(SD:<br>3.03E+04) | 5.6 x10<sup>7</sup><br>(SD:<br>3.50E+05) | 4.6 x10<sup>6</sup><br>(SD:<br>2.97E+06) | 1.3 x10<sup>5</sup><br>(SD:<br>2.32E+0) | 8.3 x10<sup>4</sup><br>(SD:<br>2.42E+08) | 4.6 x10<sup>3</sup><br>(SD:<br>1.51E+03) | 0 |
| **O4444** | 5.6 x 10<sup>8</sup><br>(SD:<br>2.34E+08) | 2.5 x10<sup>7</sup><br>(SD:<br>8.04E+07) | 1.5 x10<sup>6</sup><br>(SD:<br>4.86E+07) | 4.2 x10<sup>5</sup><br>(SD:<br>4.73E+0) | 4.1 x10<sup>4</sup><br>(SD:<br>3.50E+05) | 5.6 x10<sup>3</sup><br>(SD:<br>1.51E+04) | 0 |

Although, the viable cell count in the mature biofilms of isolates KPON2697 and KPO3875 were significantly reduced up to 2-log at sub-MIC and MIC concentrations i.e. 5mg/mL respectively, the complete biofilm eradication could be achieved using 10mg/mL NAC solution. Mature biofilms of XDR isolate KPO3337, were also markedly reduced by 2-log and 3-log reduction in biofilm cells at 5mg/mL and 10mg/mL NAC, respectively. Increasing the NAC concentration just by 2-fold (20mg/mL) resulted in the eradication of biofilms formed by isolate KPO3337 (Fig.4). A concentration dependent anti-biofilm activity of NAC was observed against other XDR-*K. pneumoniae* isolates. Although 80mg/mL NAC solution reduced the biofilm cell viability up to 6-log (99.999%), some viable cell population was still recovered. Interestingly, biofilms formed by all the isolates couldn’t tolerate 100mg/mL NAC solution and were completely eradicated after 24hrs treatment.

**Fig 4:**
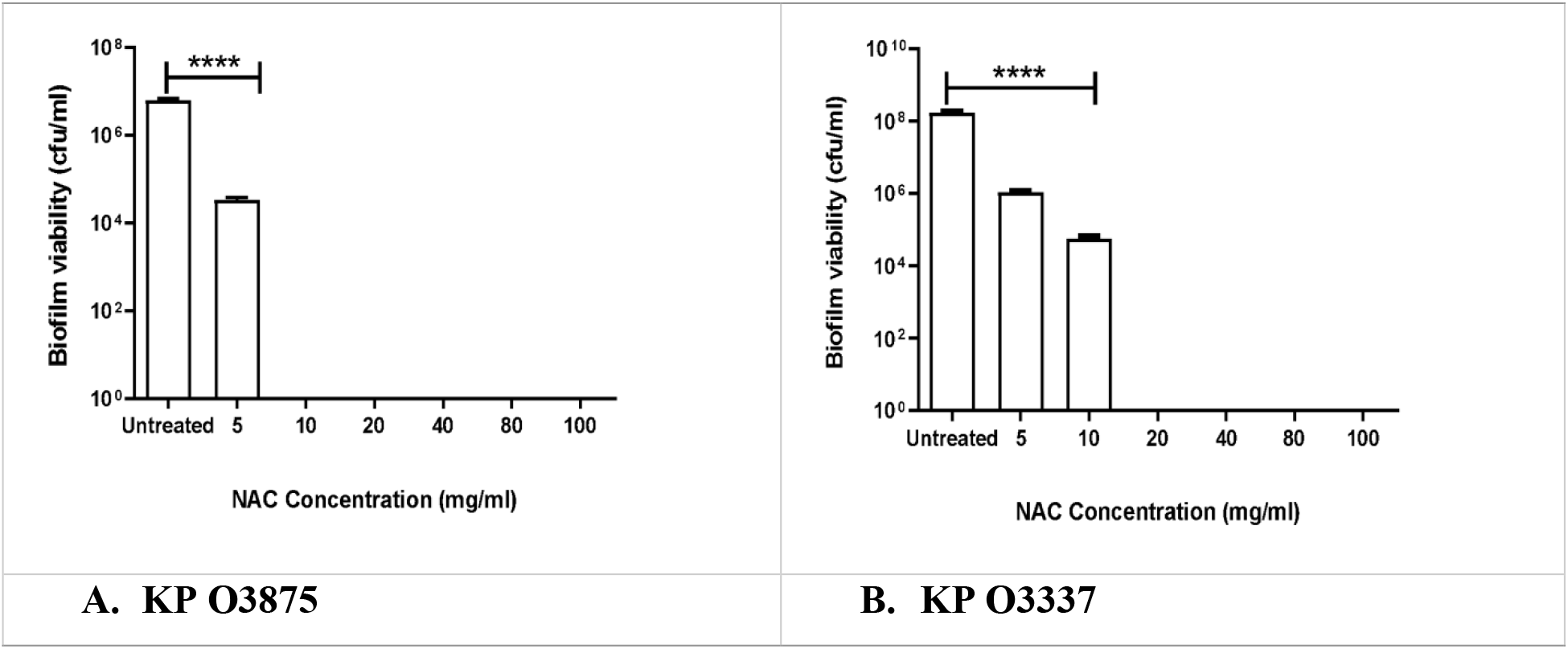
Representative graphs of *invitro* antibiofilm activity of NAC measured by CFU/ml (effect on cell viability) against XDR-*K. pneumoniae* isolates: A. KP O3875 and B. KP O3337 compared to untreated biofilm.

### 3.6. NAC is non-toxic to L929 and J774A.1 cell lines

NAC is an FDA approved compound and is considered one of the important ingredients to prepare powerful antioxidant rich food (Šalamon et al., 2019). Since NAC is an FDA approved compound and the most efficient antibiofilm concentration of NAC was 100 mg/mL that led to the total eradication of *in vitro* biofilms of XDR-*K. pneumoniae* isolates. We evaluated the cell toxicity by treating the mouse fibroblast cells (L929) and macrophage cells (J774A.1) with 80 or 100mg/mL NAC for 30mins. Incubating cells with 80 mg/mL or 100 mg/mL concentration of NAC did not exert any toxic effect on the cell lines as indicated by the high rate of cell survival of both the cell types used in the study (Fig.5). J774A.1 showed a 100% survival rate at both the tested concentrations. On the same hand, L929 also had a high survival rate of up to 91% survival at both the tested concentrations. A slight decrease in survival rate in L929 is probably due to a shock that appeared from sudden depletion in nutritional media. Higher Survival percentage in J774A.1 cell may be due to the reasons that NAC enhances the activity of macrophages (Vecchiarelli et al.,1994).

**Fig 5:**
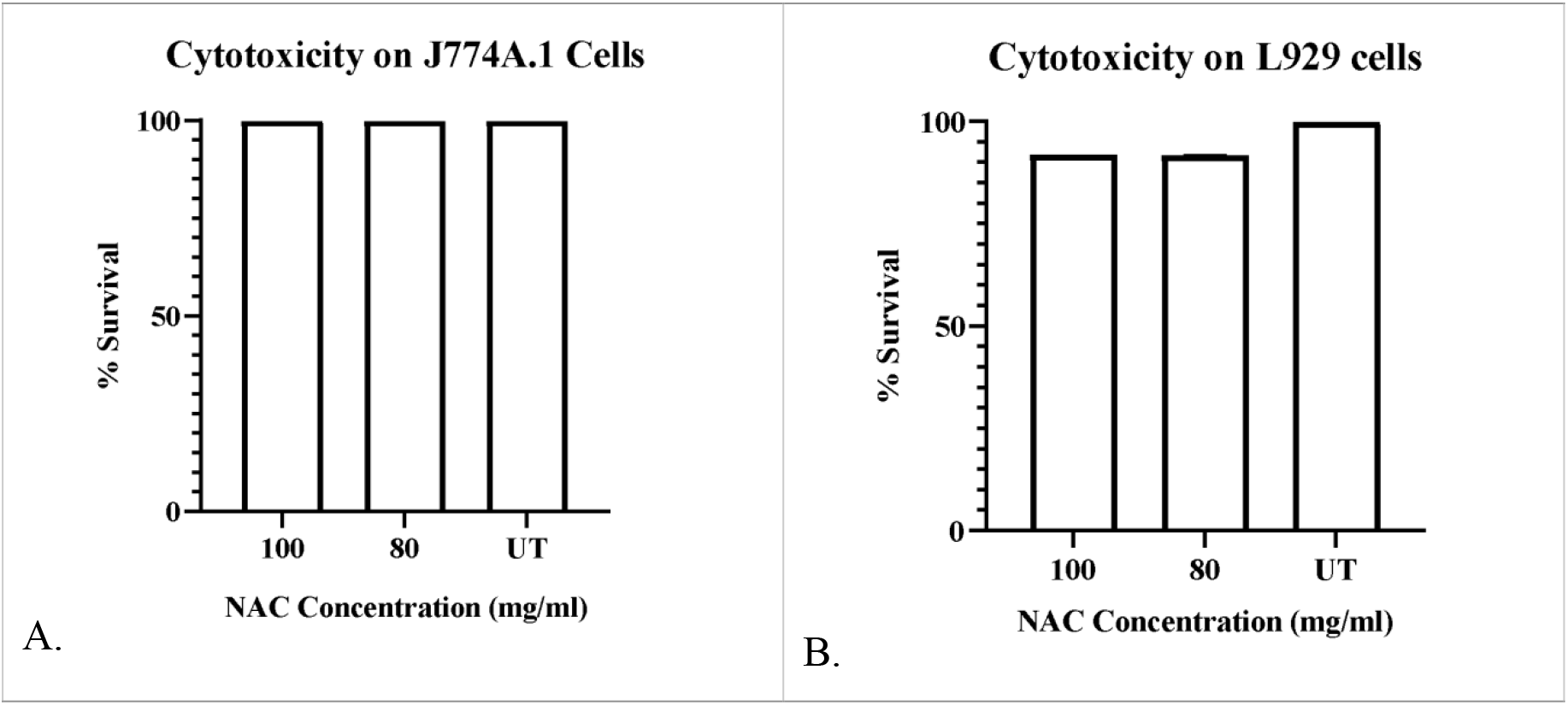
Cytotoxicity of NAC against J774A.1 and L929 cell lines after 30 mins of exposure. A) For J774A.1, 100% viability is seen post NAC treatment with both the concentrations, and B) a negligible 9% cytotoxicity was observed against L929 cells.

## 4. Discussion

*K. pneumoniae* are classified as “priority pathogens” by WHO and are one of the most important contributors to nosocomial infections (WHO 2017). The emergence of antibiotic resistance and the biofilm forming ability of pathogens including *K. pneumoniae* pose one of the greatest challenges to clinicians in treating such infections and the health system at large. Adding to the problem is a rapid rise in extensively drug resistant (XDR) bacteria that are extremely difficult to treat sometimes even due to the ineffectiveness of the last line of antibiotics. In this study, we selected XDR-*K. pneumoniae* clinical isolates to assess the efficacy of an FDA approved molecule, N-acetyl cysteine (NAC) against actively growing as well as biofilm bacteria. We report here that, of the total *Klebsiella* isolates from clinical sources, 99% were *K. pneumoniae.* Earlier clinical data also reported the high prevalence of *K. pneumoniae* species in the clinical samples (Heidary et al., 2018; Igrejas et al., 2019; Asri et al., 2021; Odari & Dawadi, 2022). Carbapenems are the first choice of clinicians for treating drug resistant -*K. pneumoniae* (Bassetti et al., 2018). Imipenem is a carbapenem antibiotic that was reported as the most effective antibiotic in treating *K. pneumoniae* associated infections. However, lately, the emergence of carbapenemase producing *K. pneumoniae* has rendered the carbapenem class of antibiotics insensitive (Papadimitriou-Olivgeris et al., 2017; Ashwath et al., 2022). In our study, we observed that imipenem resistance was predominant in up to 83% of clinical isolates. A similar scenario was also demonstrated by Ashwath et al.,2022, where more than 80% resistance among *K. pneumoniae* isolates was observed against imipenem (Ashwath et al., 2022). Besides, *K. pneumoniae* isolates also showed the least sensitivity toward other carbapenems, viz. meropenem and ertapenem (Papadimitriou-Olivgeris et al.,2017; Matovina et al., 2021., Ashwath et al., 2022)

It is worrisome to note that recent studies have shown the emergence of resistance in clinical isolates of *K. pneumoniae K. pneumoniae* against third generation cephalosporins and fluroquinolones (Xie et al.,2018; Geetha et al., 2020; Renggli et al., 2022; Rocha et al., 2022). In the present study, all the isolates of *K. pneumoniae* were resistant to cefazolin, ceftriaxone, cefotaxime, and cefuroxime. Moreover, 83% of isolates were resistant to cefoperazone sulbactam and 91% of isolates were insensitive to cefepime. Among fluoroquinolones including ciprofloxacin and ofloxacin, only 9% susceptibility was observed.

Furthermore, the rate of colistin resistance has also increased among *K. pneumoniae* (Ah et al.,2014; Zafer et al.,2019; Azam et al.,2021; Narimisa et al.,2022). In our study also we observed 25% colistin resistant isolates, which is certainly concerning. The increased antibiotic resistance is attributed to several factors, however, the prevalence of ESBL encoding genes such as *bla* _SHV_, *bla* _CTX-M_, *bla* _TEM_; and plasmids harboring quinolone resistant *qnrS, qnrA, qnrB,* and colistin resistant *mcr* genes, are regarded as the main contributor of resistance among *K. pneumoniae* (Piperaki et al., 2017; Ashwath et al., 2022).

Another crucial factor responsible for persistent *K. pneumoniae* infections is the development of biofilm on host tissues and indwelling devices. Moreover, the correlation between biofilm formation and antibiotic resistance is well established in different studies (Vuotto et al., 2017; Potempa et al., 2018; Nirwati et al.,2019). Although all the *K. pneumoniae* clinical isolates studied were capable of forming biofilms *in vitro,* 59% of the isolates were moderate biofilm formers, 33% were weak biofilm formers and only 8% were strong biofilm formers. Earlier studies also showed that clinical isolates of *K. pneumoniae* were either moderate or weak biofilm formers (Magesh et al.,2013; Cusumano et al.,2019; Singh et al., 2019). As reported by earlier studies, we also observed that all the *K. pneumoniae* isolates (25% of total *K. pneumoniae* isolates) that were resistant to all the tested antibiotics were weak biofilm formers. Similarly, we noticed that all the moderate biofilm formers are also antibiotic resistant to several antibiotics, viz. *K. pneumoniae* 100% resistance was observed against co-trimoxazole, 85% resistance was observed against imipenem, meropenem, tigecycline, and ofloxacin, and 71% resistance was observed against amoxicillin clavulanate, amikacin, gentamicin, ciprofloxacin, cefazolin, cefoperazone sulbactam, ceftriaxone, cefepime, cefotaxime, and cefuroxime. Reports suggest that the presence of plasmids carrying resistant genes in *K. pneumoniae,* modulates the expression of diverse biofilm associated genes (Ashwath et al., 2022). Possibly this may be true for the multiple drug resistant *K. pneumoniae* isolated from clinics in our study.

With the rising rates of antibiotic resistance among *K. pneumoniae,* non-antibiotic alternatives are currently being explored. N-acetyl cysteine (NAC) is an FDA approved drug used to treat acetaminophen poisoning and obstructive pulmonary diseases. Recently use of NAC as an antibacterial and antibiofilm agent has gained tremendous interest. Some studies have successfully used NAC to demonstrate its *in vitro* efficacy against planktonically grown as well as biofilms of laboratory strains of bacteria (Li et al., 2020; Aiyer et al., 2021; Chlumsky et al., 2021; Tenório et al., 2021). The antibiofilm activity of NAC is attributed to different aspects such as inhibition of EPS production, bacterial adhesion inhibition, competitive inhibition of cysteine in bacteria, interaction with bacterial proteins via thiol groups, and liquification of biofilm matrix (Zhao et al., 2010; Costa et al.,2017; Manoharan et al.,2020; Guerini et al.,2020; Guerini et al., 2022). Some reports have shown the killing efficacy of NAC is pH-mediated. NAC in its undissociated form can penetrate through the bacterial cell membrane and inside the cell, it dissociates into its conjugate base and H^+^ decreasing the intracellular pH that may interfere with DNA replication, RNA transcription, and protein translation (Kundukad et al., 2020).

In this study, we assessed the potential of NAC against the clinical isolates of XDR-*K. pneumoniae,* some of these were resistant to all the tested antibiotics. We observed that the minimum concentration of NAC required for killing the exponentially growing *K. pneumoniae* ranged between 5 to10 mg/mL and a significant number of viable cells were recovered from the cultures treated with NAC concentrations less than 5mg/mL. In 2011, Darouiche and the group also reported that the MIC and MBIC of NAC against the clinical isolates of *K. pneumoniae* were 5 and 10 mg/mL, respectively (Aslam et al., 2011).

Recently, a few groups reported that NAC was bactericidal at a concentration as low as 2mg/mL (Kundukad et al., 2020). This may be attributed to differential strain specific anti-bacterial activity of NAC and probably clinical isolates have comparatively higher tolerance toward NAC. Moreover, our cell viability results indicated that NAC is bactericidal to *K. pneumoniae* isolates contradicting previous studies that reported NAC as bacteriostatic (Manoharan et al., 2020, Aiyer et al., 2021).

However, few other studies demonstrated that NAC is bacteriostatic at low concentrations and bactericidal at higher concentrations (Perry et al., 1977; Aslam et al., 2011). We then assessed the antibiofilm activity of NAC at different concentrations against preformed 96 hr old mature biofilms of *K. pneumoniae.* Although, *in vitro* biofilms of 16.6% of isolates and of 8.3% of isolates were eradicated at 10mg/mL and 20mg/mL respectively, complete eradication of biofilms of the majority of the XDR-*K. pneumoniae* isolates could be achieved only at a minimum of 10X MIC of NAC i.e. 100mg/mL.

Biofilms are a source of recurrent infections, which is attributed to the presence of highly antibiotic-tolerant persister bacteria within biofilms, and is one of the major challenges for the management of biofilm-associated infections in the clinical setting (Darouiche RO.,2004; Lewis K., 2007; Lebeaux et al.,2014). NAC proved to be a potential therapeutic option that is able to eradicate even the persistent bacteria of XDR-*K. pneumoniae* isolates at a concentration of 100mg/mL.

Although the clinical use of acidic compounds may be potentially toxic, our choice of NAC as an alternative therapeutic to antibiotics is persuasive as several reports suggested the lack of toxicity of intravenous administration of NAC to patients in various clinical situations (Prescott et al., 1979; Smilkstein et al., 1991; Perry et al., 1998; Buckley et al., 1999).

Besides, NAC acts as a ROS scavenger and a cysteine prodrug that releases cysteine inside the cells at a slow rate helping in the restoration of the glutathione levels. (Jiao et al., 2016; Pedre et al., 2021). NAC showed cytotoxicity preventive effects on U937 monocytes and human kidney epithelial cells (HK-2 cell line) (Zhang et al., 2011; Ghani et al., 2014). Similarly, we showed that treating mouse fibroblast cells (L929) and macrophage cells (J774A.1) with NAC up to a concentration of 100mg/mL had no cytotoxicity even after 30 minutes of exposure.

In conclusion, we summarize that physiologically tolerable concentrations of NAC can completely eradicate the *in vitro* mature biofilms of all tested XDR-*K. pneumoniae* clinical isolates. Hence, we believe that the NAC deserves further preclinical and clinical assessments for effective management of WHO critical priority pathogen XDR-*K. pneumoniae* associated nosocomial infections.

## Acknowledgement

AC would like to thank UGC-BSR (F.30-487/2019(BSR)), DST-Nanomission (DST/NM/NB/2018/203), ICMR (OMI/20/2020-ECD-1) and SERB-CRG (CRG/2021/001974) for the funding support.

