## Supplementary Material for "Successfully treating biofilms of extensively drug resistant *Klebsiella pneumoniae* isolates from hospital patients with N-Acetyl Cysteine"

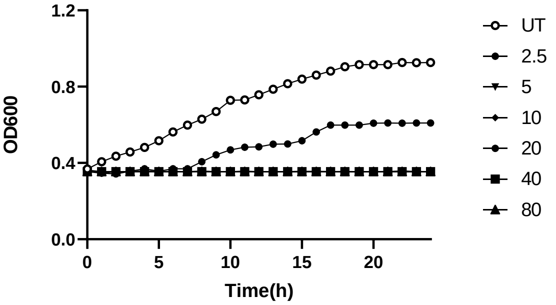


A. ON2352

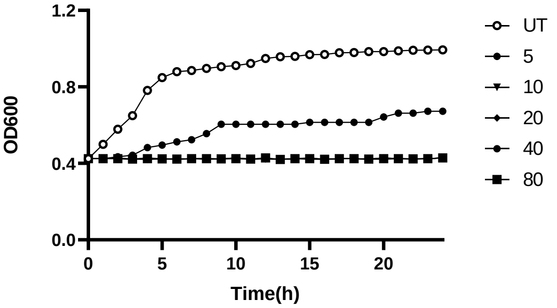


B. O3159

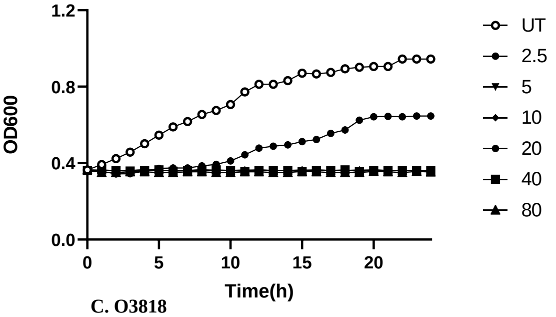


C. O3818

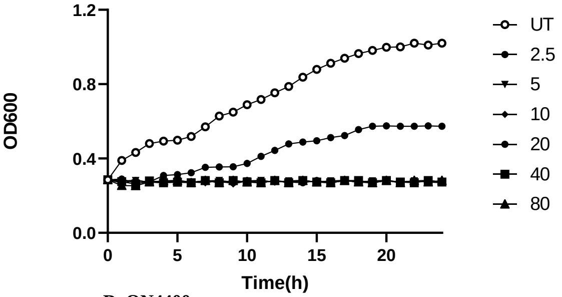


D. ON4400

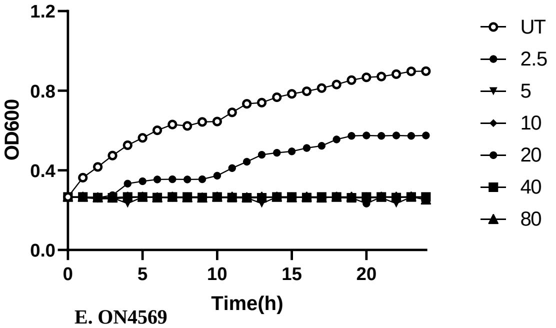


E. ON4569

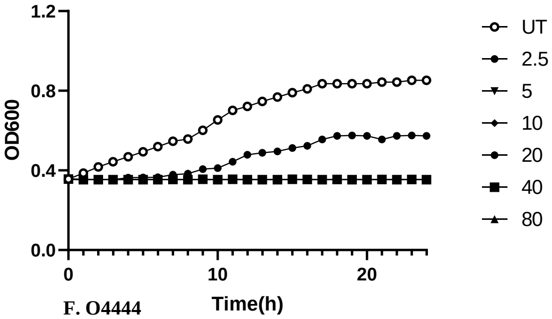


F. O4444

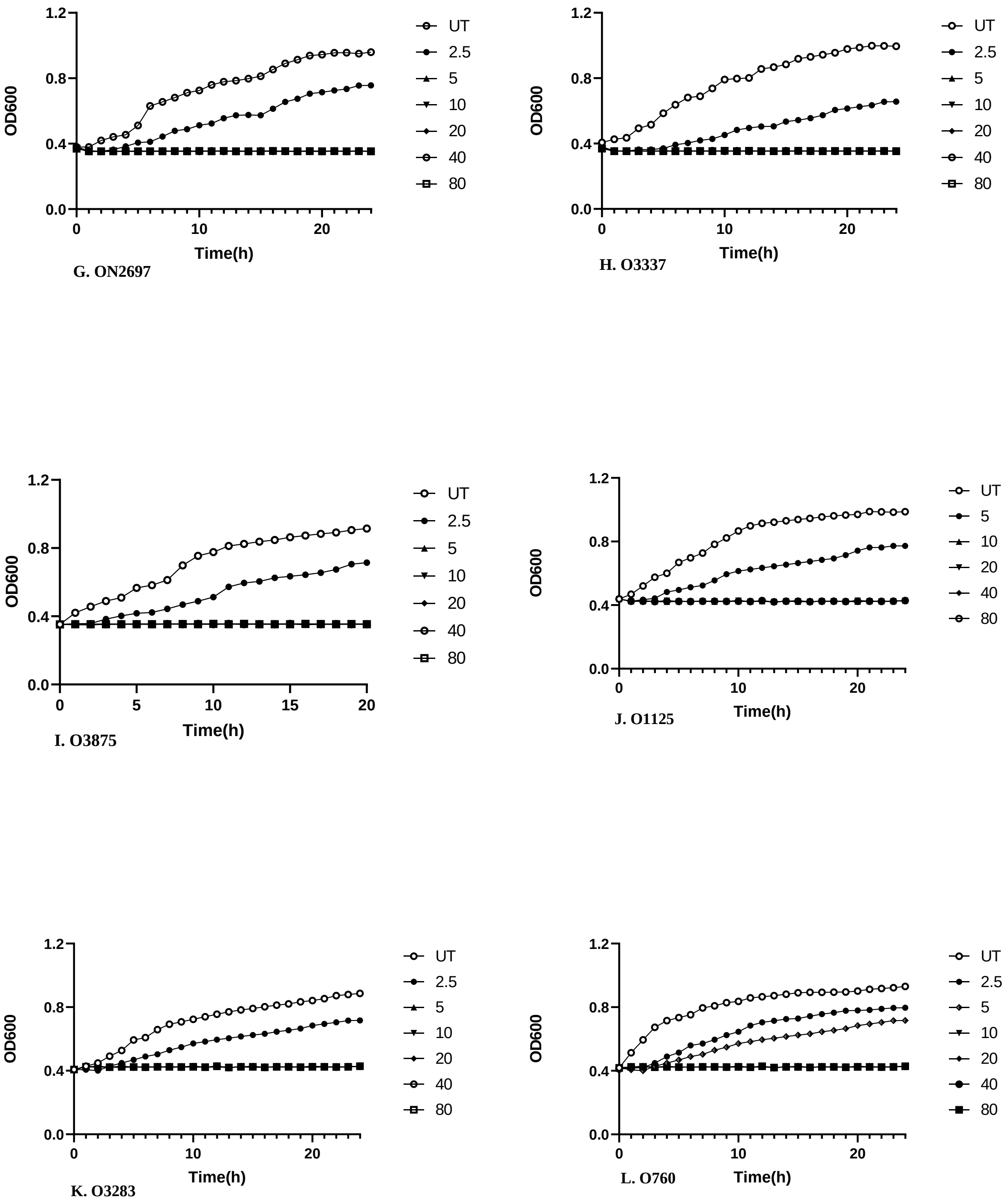


**Fig S.1 (A-L): Antibacterial activity of NAC against exponentially growing XDR-*K.pneumoniae* isolates.** All the 12 isolates were treated with different concentrations of NAC (sub-MIC to 80mg/mL) for 24h and the OD600 was measured. NAC treatment at MIC concentration and above showed complete growth inhibition among 11 isolates, except O760.
